# Autonomic activity during a daytime nap facilitates working memory improvement

**DOI:** 10.1101/2020.04.22.056580

**Authors:** Pin-Chun Chen, Lauren N. Whitehurst, Sara C. Mednick

## Abstract

Recent investigations have implicated the parasympathetic branch of the autonomic nervous system (ANS) in higher-order executive functions. These actions are purported to occur through ANS’s modulation of the prefrontal cortex, with parasympathetic activity during wake associated with working memory ability (WM). Compared with wake, sleep is a period with substantially greater parasympathetic tone. Recent work has reported that sleep may also contribute to improvement in WM. Here, we examined the role of cardiac parasympathetic activity during sleep on WM improvement in healthy young adults. Participants were tested in an operation span task (OSpan) in the morning and evening, and during the inter-test period subjects either experienced a nap or wake. We measured high frequency heart rate variability (HF HRV) as an index of cardiac, parasympathetic activity during both wake and sleep. Participants showed the expected boost in parasympathetic activity during nap, compared with wake, as well as greater WM improvement after a nap compared with an equivalent period awake. Furthermore, parasympathetic activity during sleep, but not wake, was significantly correlated with WM improvement. Together these results indicate that the natural boost in parasympathetic activity during sleep has substantial benefits to gains in prefrontal executive function in young adults. We present a conceptual model illustrating the interaction between sleep, autonomic activity, and prefrontal brain function, and highlight open research questions that will facilitate understanding of the factors that contribute to executive abilities in young adults, as well as in cognitive aging.

**Significance Statement:** Recently, the neurovisceral integration model has implicated activity on the parasympathetic branch of the autonomic nervous system (ANS) during wake in executive functioning. Parasympathetic activity peaks during deep sleep, and sleep has been shown to facilitate executive functioning. Yet, the role of parasympathetic activity during sleep for executive functioning is not known. Herein, participants demonstrated increased parasympathetic activity during deep sleep, sleep-dependent WM improvement, and associations between performance gains and parasympathetic activity in sleep, not wake. Our conceptual model illustrates the interaction between sleep, autonomic activity, and prefrontal brain function that may contribute to executive abilities in young adults and to cognitive aging.

## 1. Introduction

Working memory (WM), the ability to retain, manipulate, and update information over short periods of time for use in top-down control of complex cognitive tasks, is essential to higher order cognition and to performance of daily activities. According to the WM model by Baddeley & Hitch (Baddeley & Hitch, 1974), it comprises a verbal and visuospatial system, which are both controlled by an executive system. With these subsystems, WM has been shown to support a wide range of complex cognitive functions, including logical reasoning and problem solving, and related to measures of fluid intelligence (Conway, Cowan, Bunting, Therriault, & Minkoff, 2002; Engle, Tuholski, Laughlin, & Conway, 1999). These abilities are mediated primarily by the prefrontal cortex interacting with striatal and hippocampal areas (Miller & Cohen, 2001). Decades of work have shown strong neural activity in PFC when performing WM tasks (Funahashi, Chafee, & Goldman-Rakic, 1993; Fuster & Alexander, 1971; Levy & Goldman-Rakic, 2000). From an aging perspective, WM functions are prone to age-related cognitive decline. This decline is already evident in the normal aging process but is particularly pronounced in old-old (age > 75) adults (Hale et al., 2011). Considering the importance of WM for cognitive functions, the question of possibly modifying WM decline has been raised. Illuminating the mechanisms of WM improvement is important as this domain has been the focus of cognitive training in older adult populations with the expectation that enhanced WM will generalize to a wide range of cognitive functions and potentially slow the speed of cognitive aging.

WM training typically requires practice over a span of days, weeks or months, suggesting that offline, sleep-dependent mechanisms may be involve in the long-term improvement of WM. Sleep plays an important role in the maintenance and improvement of a wide range of cognitive processes, including the consolidation of declarative memory and procedural memory, as well as maintaining executive function, including sustained attention and WM (Könen, Dirk, & Schmiedek, 2015; Vriend et al., 2013). Indeed, it’s known that sleep deprivation/restriction detrimentally affects WM. For example, sleep deprivation/restriction lead to impairment in sustained attention (Goel, Rao, Durmer, & Dinges, 2009; Lo et al., 2012) and a variety of cognitive tasks involving WM, such as digit span. (Quigley, Green, Morgan, Idzikowski, & King, 2000) and N-back tasks (Choo, Lee, Venkatraman, Sheu, & Chee, 2005), effects likely driven, in part, by altered functioning of frontal and parietal networks (Chee & Choo, 2004). Although studies have repeatedly demonstrated that a sleep-deprived brain, compared with a well-rested one, performs worse on WM tasks (Lo et al., 2012; 2016), much less is known about the direct contribution of sleep-specific mechanisms supporting WM improvement. Recently, studies that directly tested the effect of post-training sleep on WM performance suggested that a period of sleep, compared to wake, facilitates WM. (Zinke, Noack, & Born, 2018; Kuriyama, Mishima, Suzuki, Aritake, & Uchiyama, 2008a; Lau, Wong, Lau, Hui, & Tseng, 2015). In these studies, training adult participants on an N-back task over several sessions improved accuracy of performance, but only if the interval between training sessions included nocturnal sleep (Zinke et al., 2018; Kuriyama et al., 2008a) or a nap (Lau et al., 2015), in comparison with daytime periods of wakefulness. One plausible neurophysiological mechanism for such training improvement is slow wave sleep (SWS). SWS has received increasing attention due to its roles in offline memory consolidation and memory reactivation (Berkers et al., 2018; Marshall & Born, 2007). In addition, SWS has been linked to synaptic plasticity and cortical reorganization (Tononi and Cirelli, 2003; Takashima et al., 2006; Dang-Vu et al., 2010). Intriguingly, several studies have shown a specific association between electrophysiological (EEG) activity during SWS in the enhancement of WM. Pugin and colleagues (2015) demonstrated a correlation between slow wave activity (SWA) during SWS in frontal areas and WM performance after three weeks of WM training (Pugin et al., 2015). Furthermore, SWA during SWS has been shown to predicted WM gains across a period of sleep in both young (Ferrarelli et al., 2019) and older adults (Sattari et al., 2019). Taken together, these studies suggest that SWS might provide an optimal brain state for the improvement of WM.

A different line of research has demonstrated a significant contribution of the autonomic nervous system (ANS) for WM. Cardiac vagal tone, which represents the contribution of the parasympathetic nervous system to cardiac regulation is known for its role in regulating involuntary bodily functions, such as breathing, heart rate and digestion, and is less recognized for its role in influencing cognitive processing. Yet, over recent decades, vagal activity is acknowledged to be linked with self-regulation at the cognitive, emotional, social, and health levels. Thayer and others have published a body of research implicating cardiac vagal influence on a range of cognitive abilities supported by the prefrontal cortex (PFC) (Lane, Reiman, Ahern, & Thayer, 2001; Smith, Thayer, Khalsa, & Lane, 2017; Thayer & Lane, 2009). Descending projections from the PFC to the brainstem and hypothalamic structures allow for bi-directional communication between the central nervous system and the ANS through the vagus nerve (Packard et al., 1995; Thayer et al., 2009), and thus prominent models of ANS and cognition, including the Neurovisceral Integration Model (Smith et al., 2017; Thayer & Lane, 2009), have focused on the impact of vagal cardiac activity on executive function cognition. In humans, a well-established method to non-invasively examine autonomic activity is heart rate variability (HRV), which measures systematic variation in the beat-to-beat interval (Shaffer et al., 2014). The most commonly used HRV analytical approaches are time domain analysis and frequency domain analysis. The primary time domain measure is the root mean square of successive differences (RMSSD), which reflects the beat-to-beat variance in heart rate and is used to estimate vagally-mediated changes in the RR time series (Laborde, Mosley, & Thayer, 2017; Shaffer et al., 2014). For frequency domain analysis, spectral analysis of the cardiac signal in the high frequency range (HF HRV: 0.15-0.40 Hz) is indicative of vagally-mediated respiration and parasympathetic cardiac activity. Vagally-mediated HRV (e.g. RMSSD, HF HRV) during wake has been shown to predict performance on a wide range of cognitive tasks that rely on PFC activity (Thayer et al., 2009). For example, compared to individuals with low resting vagally-mediated HRV, high HRV individuals perform better on both WM (n-back task: Hansen et al., 2003; operation-span task: Mosley et al., 2018) and cognitive inhibition (i.e. Stroop task; Hansen et al. 2004). Additionally, reducing HRV, via aerobic de-training, comes at significant cost to executive functioning (Hansen et al., 2004). More recently, studies have demonstrated that directly stimulating the vagus nerve can increase vagally-mediated HRV (Clancy et al., 2014), improve verbal memory (Clark et al., 1999; Jacobs et al., 2015), and accelerate extinction learning (Burger et al., 2016). These studies suggest that strong modulation of ANS activity may benefit prefrontal functioning.

Sleep strongly modulates ANS activity (Baharav et al., 1995). As the brain shifts from wake into sleep, the body also undergoes marked changes with heart rate deceleration and relative increases in parasympathetic HF HRV across the three stages of non-rapid eye movement (NREM) sleep (i.e., Stage 1, Stage 2, and Slow Wave Sleep (SWS)) (Trinder et al., 2001). Additionally, similar HRV profiles have been shown between daytime (naps) and nighttime sleep (Cellini, Whitehurst, McDevitt, & Mednick, 2016; Whitehurst, Naji, & Mednick, 2018). It is not known whether naturally elevated vagal activity during NREM sleep might support WM.

We investigated the impact of parasympathetic activity during sleep versus wake on both general WM performance (baseline at Test 1) and WM improvement across a day (difference score between Test 2-Test 1). We examined WM using the Operation Span Task (OSpan), which is a dual-task consisting of a processing subtask and a short-term memory subtask that has been commonly used to test central constructs of WM, but has not been examined in the context of sleep. Thus, the current study aimed: (1) to assess the cardiac activity across sleep stages during a daytime nap in healthy young adults; (2) to compare the effect of a daytime nap versus wake on WM improvement; and (3) to explore the impact of parasympathetic activity during sleep and wake on WM. We hypothesized that participants would show increases in parasympathetic activity during NREM sleep compared to waking and REM sleep. Furthermore, we predicted that sleep, especially SWS, would benefit WM to a greater extent than wake, and that parasympathetic activity during SWS would be positively associated with WM improvement to a greater extent than waking activity.

## 2. Material and methods

### 2.1. Participants

104 young adults (Age:17-23 [Mean=20.7, SD= 2.95], 60 males) with no personal history of neurological, psychological, or other chronic illness provided informed consent, which was approved by the University of California, Riverside Human Research Review Board. For participants under the age of 18 years, informed consent was obtained from a parent and/or legal guardian. All methods were performed in accordance with the relevant guidelines and regulations. Participants were randomized to either have a 2-hour nap opportunity monitored with polysomnography (PSG) (Nap, n=53), or stay awake (Wake, n=51), where subjects engaged normal daily activities with activity watch monitoring, but were not allowed to have caffeine or take a nap. Participants included in the study had a regular sleep-wake schedule (reporting a habitual time in bed of about 7–9 h per night), and no presence or history of sleep, psychiatric, cardiovascular, or neurological disorder determined during an in-person, online, or telephone interview. Participants received monetary compensation for participating in the study.

### 2.2. Working memory task

The current study used the Operation Span Task (OSpan)^66^ as a measure of WM capacity, which requires participants to solve a series of math operations while memorizing a set of unrelated letters. The task was programmed in Matlab (The MathWorks Inc., 2015) using Psychtoolbox, which allows random generation of stimuli every trial. The task included 3 practice and 40 test trials. Participants were tested in letter strings four and seven. For each letter string, participants were shown a series of math problems that they had to confirm were correct within 3 seconds, using pre-determined responses on the keyboard. After each equation, a letter would appear on the screen and the subject was instructed to remember each letter. At the end of each string, the participant was instructed to recall the letters in the order presented by typing responses on a computer keyboard. Immediately after each trial, the next letter string would be presented. An example of a four-item trial might be: 12 − 2 = 8 (correct/ incorrect?) → J; 6 + 7 = 14 (correct/incorrect?) → G; 3 − 2 = 1 (correct/incorrect?) → S; 5 + 7 = 13 (correct/incorrect?) K (see Figure 1b). After verifying the four equations in this example, participants were asked to type the presented letters in the order that they were presented (in this case JGSK). If the participants forgot one of the letters in a trial, they were instructed to provide their best guess. In addition, to decrease trade-off between solving the operations and remembering the letters, a 70% accuracy criterion on the math operations was required for all the participants. We excluded 1 participant in the Nap group based on this criterion. We calculated performance as: number of correct letters recalled/ total number of letters in the string per trial, and then we averaged over the total 40 trials. For assessing change in performance from session 1 and session 2, we calculated the difference in performance between the two sessions (session 2 – session 1).

**Figure 1.**
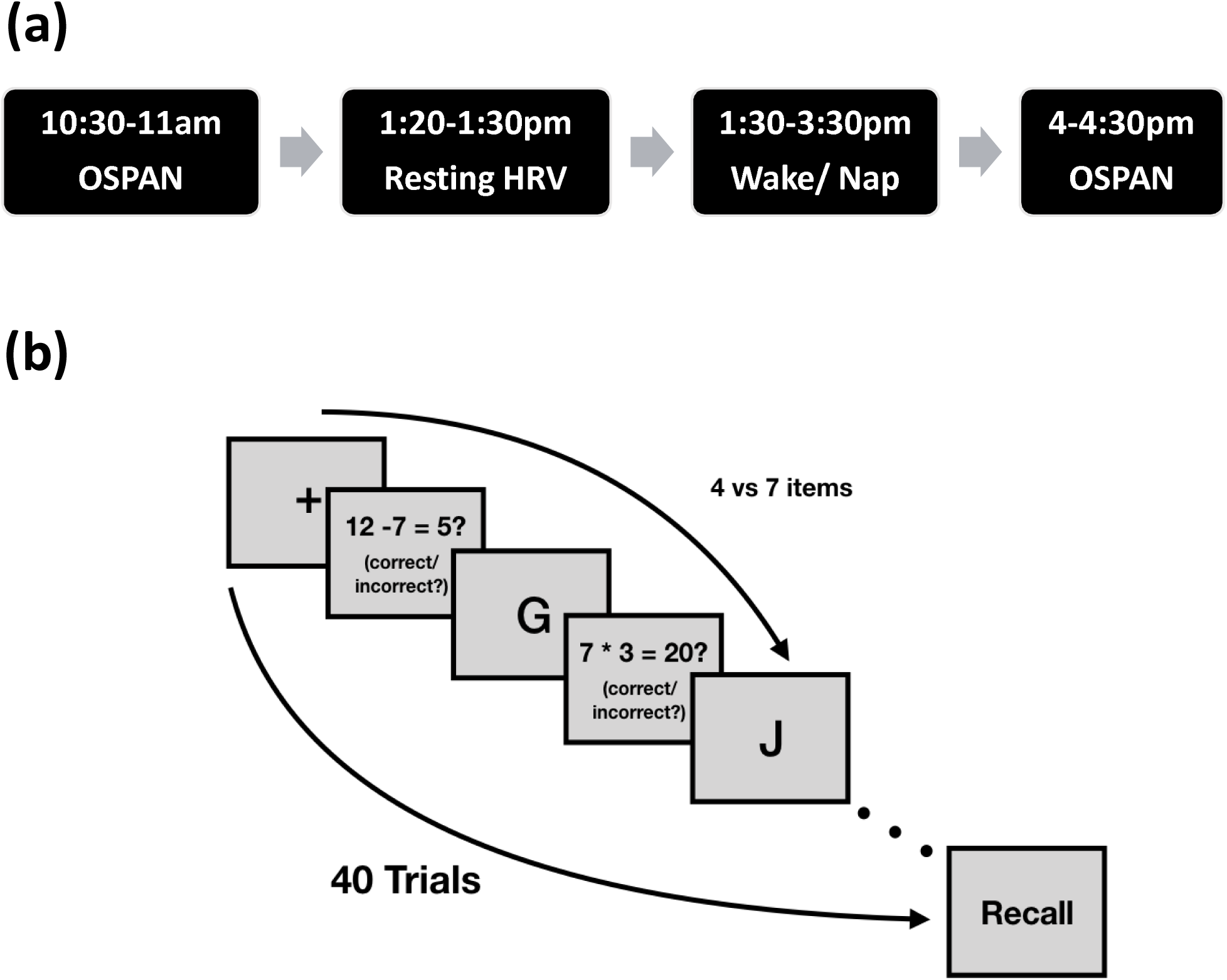
Study design and WM task (OSpan) (a) Subjects completed the 30-min OSpan WM task at 10:30 AM. Next, they were randomly assigned to either a nap condition (Nap) or a wake condition (Wake) that took place between 1:30-3:30PM. Between 4 to 4:30 PM, subjects repeated the WM task. (b) The OSpan task included 3 practice and 40 test trials. Participants were tested in letter strings four and seven. For each letter string, participants were shown a series of math problems that they had to confirm were correct within 3 seconds by entering responses on the keyboard. After each equation, a letter would appear on the screen and the subject was instructed to remember each letter. At the end of each string, the participant was instructed to recall the letters in the order presented by typing responses on the keyboard. Immediately after each trial, the next letter string would be presented.

### 2.3. Study Procedure

Participants were asked to refrain from consuming caffeine, alcohol, and all stimulants for 24 h prior to and including the study day. Participants filled out sleep diaries for one week prior to the experiment and wore wrist-based activity monitors the night before the study (Actiwatch Spectrum, Philips Respironics, Bend, OR, USA) to ensure participants were well-rested (at least 7 hours per night for the youngers and 6 hours for the elders during the week including the eve of the experimental day). On the experimental day (Figure 1a), participants arrived at the Sleep and Cognition lab at 10:00AM and had EEG electrodes applied, followed by an Operation Span (OSpan) WM task. Nap/wake interventions occurred between 1:30-3:30 PM. At 1:30PM, Nap subjects took a polysomnographically-recorded nap and were given 2-hours time-in-bed to obtain up to 90-min total sleep time. Sleep was monitored online by a trained sleep technician. In the wake group, subjects were asked not to nap, exercise, or consume caffeine or alcohol, and were monitored with actigraphy during the break. In addition, the EEG/ECG was not recorded during wake break. Between 4 and 4:30PM, all subjects were retested on the memory task. Subjects completed the Karolinska Sleepiness Scale (KSS; T Åkerstedt, 1990) questionnaire two times throughout the experimental day; at the start of each WM task (Session 1 and Session 2) to report their sleepiness. KSS is a 9-point Likert scale often used when conducting studies involving self-reported, subjective assessment of an individual’s level of drowsiness at the time, in which a higher score yields a sleepier state at that time.

### 2.4. Sleep Recording and Scoring

EEG data were acquired using a 32-channel cap (EASYCAP GmbH) with Ag/AgCI electrodes placed according to the international 10-20 System^31^. 22 out of 32 electrodes were active scalp recordings. The remaining electrodes were used for electrocardiogram (ECG), electromyogram (EMG), electrooculogram (EOG), ground, an online common reference channel (at FCz location, retained after re-referencing), and mastoid (A1 & A2) recordings. The EEG was recorded with a 1000 Hz sampling rate, amplified (ActiCHamp), and was re-referenced to the contralateral mastoid (A1 & A2) post-recording. Only eight scalp electrodes (F3, F4, C3, C4, P3, P4, O1, O2), the EMG and EOG were used in the scoring of the nighttime sleep data. High pass filters were set at. 3 Hz and low pass filters at 35 Hz for EEG, EOG and EMG. Raw data were visually scored in 30-sec epochs into Wake, Stage 1, Stage 2, Slow Wave Sleep (SWS; Stages 3 and 4) and rapid eye movement sleep (REM) according to the Rechtschaffen & Kales’ manual using HUME, a custom MATLAB toolbox. Prior to sleep scoring, data were pre-processed using BrainVision Analyzer 2.0 (BrainProducts, Munich Germany). Minutes in each sleep stage were calculated and sleep latency (SOL) were calculated as the number of minutes from lights out until the initial epoch of sleep, Stage 2, SWS and REM. Additionally, wake after sleep onset (WASO) was calculated as total minutes awake after the initial epoch of sleep and sleep efficiency (SE) was computed as total time spent asleep after lights out divided by the total time spent in bed x 100. Durations in each stage were shown in Table 1.

**Table 1.** Descriptive statistics for nap

|  | Mean | SEM |
| --- | --- | --- |
| TIB (min) | 78.069 | 3.1 |
| TST (min) | 61.500 | 2.94 |
| SOL | 7.664 | 0.99 |
| Stage 1 | 4.586 | 0.47 |
| Stage 2 | 31.448 | 1.94 |
| SWS | 19.224 | 1.96 |
| REM | 6.241 | 1.09 |
| WASO | 6.560 | 0.88 |
| SE (%) | 77.588 | 2.21 |

Note: SEM = Standard Error Mean; TIB = Total Time in Bed; TST = Total Sleep Time; SWS = Slow Wave Sleep; REM = Rapid Eye Movement sleep; WASO = Wake After Sleep Onset (calculated as the minutes of wake after first epoch of sleep); SOL = Sleep Latency (calculated as the time to first epoch of sleep); SE = Sleep Efficiency (calculated as $100 \times \text{TST} / \text{TIB}$ ). All measures are represented in minutes besides SE which is in percentage.

### 2.5. Heart Rate Variability

Electrocardiogram (ECG) data were acquired at a 1000-Hz sampling rate using a modified Lead II Einthoven configuration. We analyzed HRV of the R-waves series across the whole sleep/wake period using Kubios HRV Analysis Software 2.2 (Biosignal Analysis and Medical Imaging Group, University of Kuopio, Finland), according to the Task Force guidelines (Malik et al., 1996). RR peaks were automatically detected by the Kubios software and visually examined by trained technicians. Incorrectly detected R-peaks were manually edited. Missing beats were corrected via cubic spline interpolation. Artifacts were removed using the automatic medium filter provided by the Kubios software. The HRV analysis of the RR series was performed by using an independent lab tool. An autoregressive model (Model order set at 16) (Boardman, Schlindwein, Rocha, & Leite, 2002) was employed to quantify the absolute spectral power (ms^2^) in the LF HRV (0.04–0.15 Hz; ms^2^) and the HF HRV (0.15–0.40 Hz; ms^2^) frequency bands. The LF HRV and HF HRV measures had skewed distributions and as such were transformed by taking the natural logarithm, as suggested by Laborde et al. (2017). From these variables we derived the HF normalized units (HF_nu_= (HF HRV[ms^2^]/ HF HRV[ms^2^] + LF HRV[ms^2^])*100). Since the LF normalized units are mathematically reciprocal to HF_nu_ (i.e. LF_nu_ =1-HF_nu_), to avoid redundancy, we computed only the HF_nu_ index, an index often thought to reflect vagal modulation^8^. Besides frequency domain, we also calculated a time domain measure typically used to assess parasympathetic activity, RMSSD. This value is obtained by first calculating each successive time difference between RR intervals in milliseconds. Then, each of the values is squared and the result is averaged before the square root of the total is obtained. Similar to the frequency adjustments, to adjust for the unequal variance in the RMSSD, we report the natural logarithm of RMSSD. Additionally, we included the RR interval as an index of cardiac autonomic control in our analyses.

For the analysis of RR, HR and frequency-domain HRV measures during different sleep stages, consecutive, artifact-free windows of undisturbed sleep were selected across the nap. Each window was 3-min in duration and the 1.5-min preceding and the entire 3-min epoch were free from stage transitions and movement times. Windows were identified and averaged within Stage 2, SWS and REM sleep. We also analyzed 3 min of pre-nap wakefulness (Wake). Epochs of stage 1 and wake after sleep onset were not analyzed, because these periods have not been previously reported to contribute to memory and are hard to isolate in the recording. This methodology emphasizes consolidated sleep stages and because naps have more fragmented sleep due to increased stage transitions, this method of HRV analysis decreased the number of subjects that could be analyzed.

### 2.6. Data Reductions

104 (Males = 60) young adults were recruited and randomized into three nap conditions (Wake=51, Nap=53). 1 participant were excluded based on Math accuracy (70%). Therefore, for the WM task, we have 103 (Wake=51, Nap=52) participants in our dataset. For ANS measures, 5 participants nap recordings were not collected due to recording computer failures. For Stage 2 sleep, we excluded 6 participants due to no 3-minute window of undisturbed consecutive Stage 2 sleep. For SWS sleep, we excluded 14 participants due to no 3-minute window of undisturbed consecutive SWS. For REM sleep, we excluded 30 participants due to no 3-minute window of undisturbed consecutive REM sleep. In summary, 47 participants were included in Wake; 41 participants were included in Stage 2; 33 participants were included in SWS; 16 participants were included in REM sleep.

### 2.7. Statistical Analyses

In order to investigate within-subject profile of cardiac activity across sleep stages, we used a linear-mixed effect models (LME), which do not depend on limited assumptions about either variance-covariance matrix assumptions (sphericity) or complete data. As the numbers of subjects are different among different sleep stages, LME corrects degrees of freedom with Satterthwaite approximation. Our LME model used a within-subjects factor of stage (Wake, Stage 2, SWS, REM). All comparisons were adjusted by Bonferroni correction.

To confirm that there was no difference in WM baseline performance between the two Nap conditions, we used a one-way analysis of variance (ANOVA) with Nap Condition (Wake, Nap) as the between-subject factor, Test 1 as the dependent variable. To test the difference in WM change across the day, we used a one-way ANOVA with Nap Condition (Wake, Nap) as the between-subject factor, Test 2 – Test 1 as the dependent variable. To examine whether sleepiness level changed with different nap conditions, we used a repeated-measure ANOVA with Nap Condition (Wake, Nap) as the between-subject factor, Session (1 and 2), as the within-subject factor, and KSS as the dependent variable. Pearson correlation coefficients were used to examine the bivariate relationship between HRV variables of interests and WM performance measures, relationship between sleep parameters and WM performance, as well as relationship between KSS and WM performance.

To assess the relative importance of HRV variables for WM improvement, we utilized a hierarchical, linear regression approach. In Model 1, baseline WM performance was the independent variable and Test 2 was the dependent variable. In Model 2, we added the HRV factors as independent variables. By comparing Model 1 and 2, we measure the explanatory gain of HRV factors over and above individual differences in WM baseline performance. To compare between two nested models, we conducted a likelihood ratio test. Under the null hypothesis, the full/ restricted model (Model 2) is just as good as the reduced/ unrestricted model (Model 1). Therefore, a significant result on this test indicated that overall model fit is improved after adding the predictors in Model 2.

## 3. Results

### 3.1. HRV during Wake and Sleep

Descriptive statistics for sleep architecture were shown in Table 1. Descriptive statistics for autonomic profiles across sleep stages were shown in Table 2.

**Table 2.** Summary of HRV Parameters Across Sleep Stages

| Stage | Wake | Stage 2 | SWS | REM |
| --- | --- | --- | --- | --- |
| RR (ms) | 949.46 | 1006.61 | 1001.21 | 923.71 |
|  | (21.76) | (25.51) | (30.73) | (40.28) |
| RMSSD | 4.24 | 4.35 | 4.23 | 4.14 |
|  | (0.08) | (0.07) | (0.11) | (0.17) |
| HF HRV | 6.77 | 7.04 | 6.87 | 6.52 |
|  | (0.15) | (0.14) | (0.19) | (0.32) |
| HFnu | 52 | 59 | 71.8 | 46.6 |
|  | (2.3) | (2.5) | (2.7) | (3.5) |

Data are reported as Mean. Standard errors are shown in the rows below the Mean. Note: RR = RR intervals; RMSSD = Root mean square of successive differences; HF HRV= Power in the Low Frequency band of the HRV spectrum, often between 0.04 - 0.15 Hz; HFnu = HF/(HF+LF)%; N2 = Stage 2 Sleep; SWS = Slow Wave Sleep; REM = Rapid Eye Movement sleep.

Prior studies have reported increasing parasympathetic activity from waking to deeper stages of NREM sleep. In order to test this autonomic profile in each age group, we used an LME model examining HRV variables across sleep stages (Wake, Stage 2, SWS, REM). We found a stage effect for RR intervals (Figure 2a) [F_(3,33)_ = 15.598, p < 0.001], with a significant lengthening of RR intervals during Stage 2 [p < 0.001] and SWS [p= 0.001] relative to Wake. Similarly, RMSSD showed changes across sleep stages (Figure 2b) [F_(3,33)_ = 6.092, p = 0.002], with a significant higher heart rate variability during Stage 2 [p < 0.001], and a non-significant difference in SWS [p = 0.88] relative to Wake. The HF HRV showed a stage effect (Figure 2c) [F_(3,33)_ = 10.912, p < 0.001], with a higher but not significant vagal tone during Stage 2 [p = 0.014] and SWS [p = 0.74], relative to Wake. HF_nu_ showed a stage effect [F_(3,33)_ = 28.404, p < 0.001], with a marked increase of vagal tone in SWS (p < 0.001), compared with Wake. During REM sleep, participants showed significantly shorter RR intervals compared to Stage 2 [p = 0.001] and SWS [p = 0.002], as well as significantly lower HF_nu_ compared to Stage 2 [p < 0.001] and SWS [p < 0.001].

**Figure 2.**
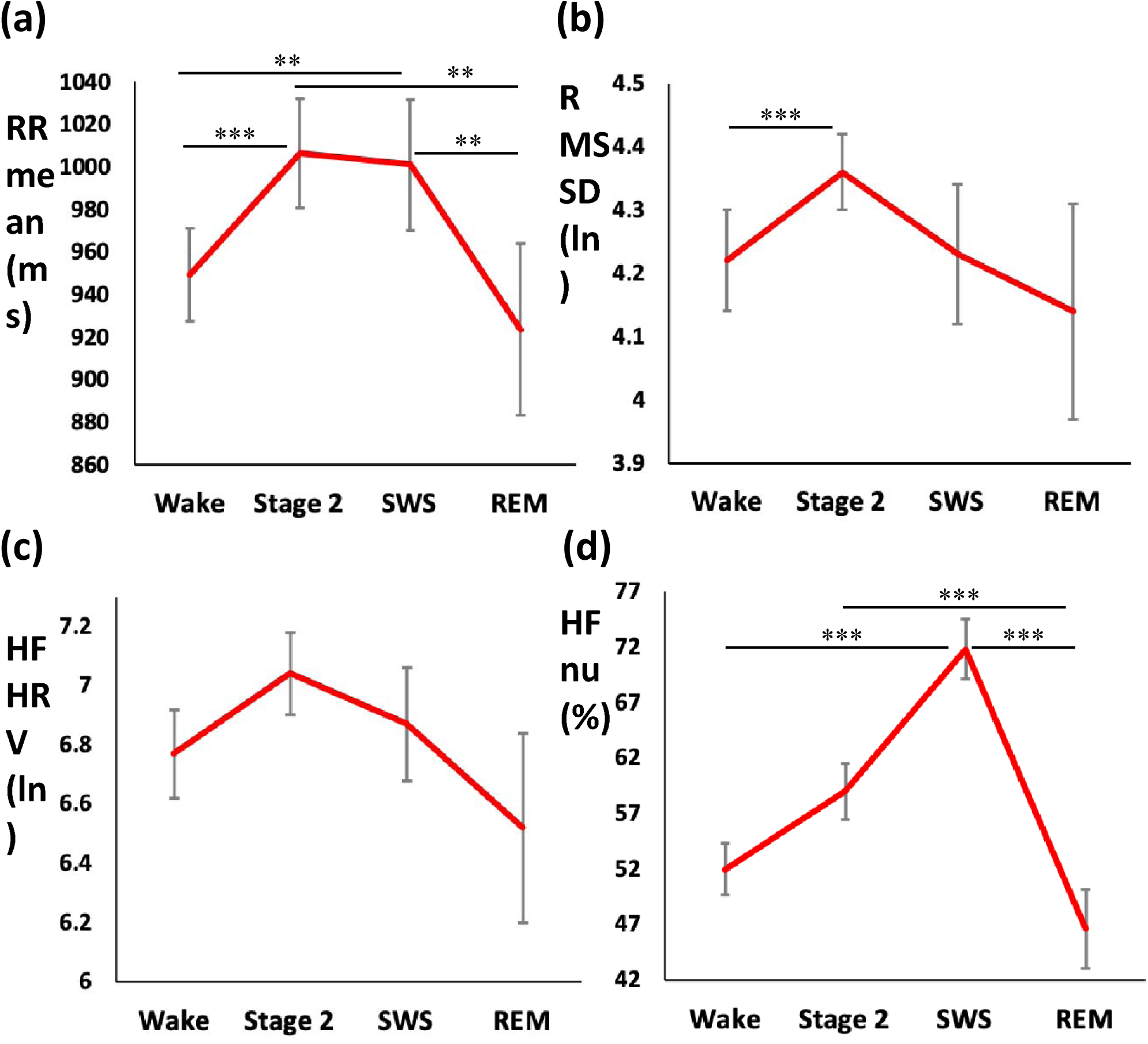
(a–d) Heart rate variability (HRV) components across sleep stages. (a) Mean of RR intervals (ms) (b) RMSSD (ln) (c) HF HRV (ln) (d) HFnu (%). Asterisks above bars indicate significant differences between stages (*p < 0.05; **p < 0.01; ***p < 0.001). Error bars represent standard error of the mean. The between-stage effects were based on the LME model examining HRV variables across sleep stages (Wake, Stage 2, SWS, REM).

### 3.2. Working Memory Performance: Comparing Nap vs Wake Group

Our analysis revealed no significant difference in WM between the two nap conditions at baseline [F(1,101) = 0.79, p = 0.376]. We compared differences in WM improvement after either a nap or wake period using a one-way ANOVA with Nap condition (Nap vs. Wake) as the independent variables, Test 2-Tset 1 WM performance as the dependent variable. The analysis revealed a main effect of nap condition [F(1,101) = 3.992, p = 0.048], in which participants showed a greater differential benefit from the nap compared to the wake condition.

**Figure 3.**
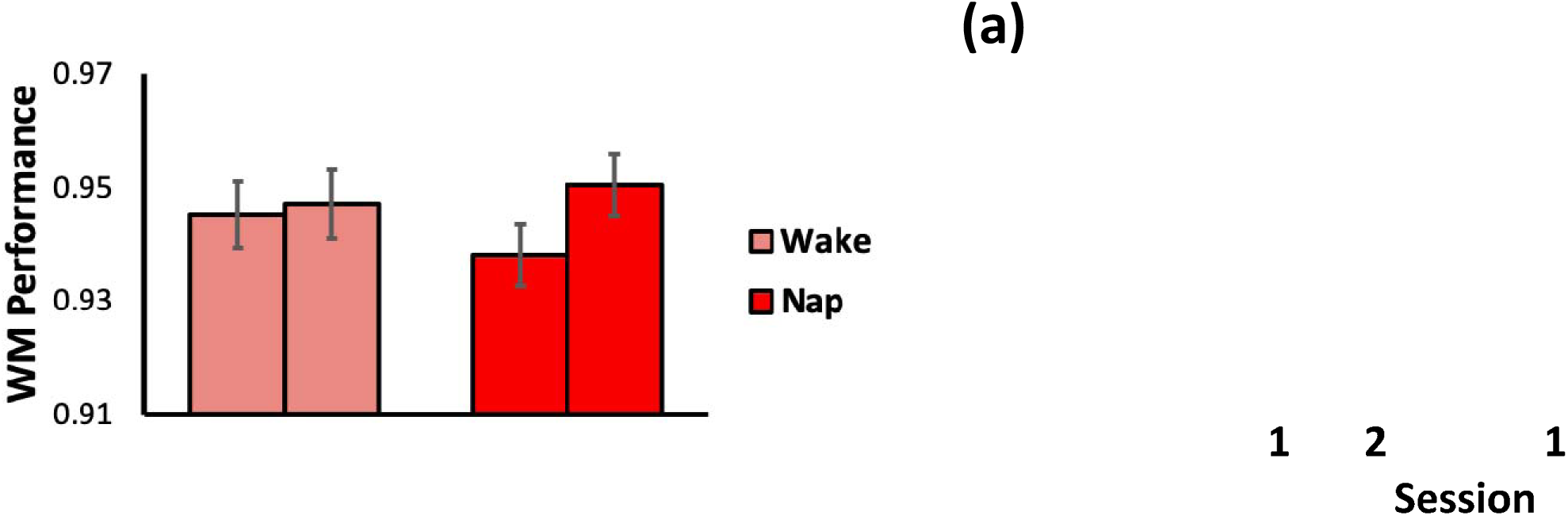
(a–b) Working Memory Performance by Nap Condition: (a) Session 1 and 2 performances by Nap Condition (b) WM improvement by Nap Condition: Significant difference in WM improvement after a nap or a period of wake was observed. Asterisks between error bars indicate significant differences between nap conditions (*p < 0.05). Error bars represent standard error of the mean.

Repeated-measure revealed no main effect of session or nap condition on sleepiness (KSS), but a significant interaction between session and nap condition [F_(1,99)_ = 5.445, p = 0.021], where the sleepiness level significantly decreased after the nap. Neither the morning nor the afternoon KSS measures were correlated with WM performances [all ps > 0.23]. Descriptive statistics for KSS were shown in Table 3.

**Table 3.** Karolinska Sleepiness Scale (KSS) scores across the day

| Session | Wake (N=51) | Nap (N=54) |
| --- | --- | --- |
| Session 1 | 5.31 (0.29) | 5.96 (0.28) |
| Session 2 | 4.98 (0.30) | 3.35 (0.20) |

*Data are reported as Mean (Standard Error).*

### 3.3. Associations between Parasympathetic Activity during Wake and Sleep on Working Memory

Next, we examined the impact of parasympathetic activity during wake and sleep on WM performance and improvement. We used Pearson correlation coefficients to examine the relationship between parasympathetic activity as measured by HF^nu^, HF HRV (ln), and RMSSD (ln) and WM performance (baseline and improvement). WM baseline performance was not correlated with HRV measures during stage 2 sleep (all ps > 0.150), SWS (all ps > 0.184), REM sleep (all ps > 0.092), or Wake (all ps > 0.439). In alignment with our expectation, WM improvement was positively correlated with SWS HFnu (r = 0.449, p = 0.015, Figure 4a). Similar positive, marginally-significant associations were also found between WM improvement and HF HRV (ln) as well as RMSSD (ln) during SWS (HF HRV: r = 0.367, p = 0.05, Figure 4b; RMSSD: r = 0.347, p = 0.065, Figure 4c). However, there was no significant associations between WM improvement and autonomic activities during stage 2 sleep (all ps > 0.434), REM sleep (all ps > 0.180), or Wake (all ps > 0.584). Furthermore, sleep alone (total time in bed, total sleep time and time in each stage (minutes)) was not significantly correlated with WM baseline or improvement (all ps > 0.41).

**Figure 4.**
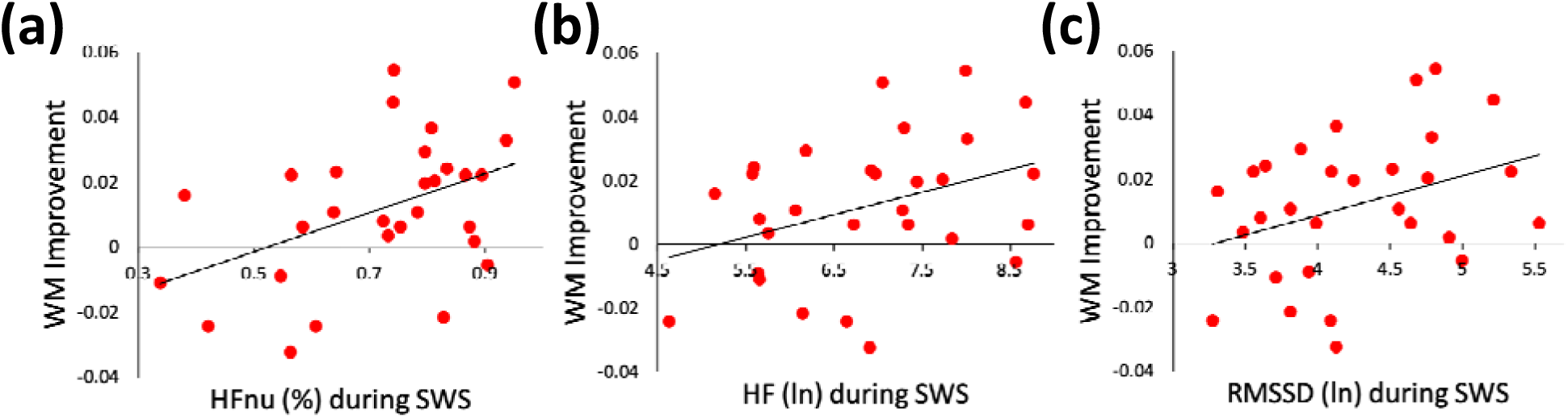
Working Memory improvement and Autonomic Activity. Association between WM improvement and (a) HFnu (%) during SWS (r = 0.449, p = 0.015); and (b) HF HRV (ln) during SWS (r = 0.367, p = 0.05); (c) RMSSD (ln) during SWS (r = 0.347, p = 0.065).

Next, we assessed the importance for memory performance of ANS activity using hierarchical, linear regressions. Two linear regression models were built to predict WM session 2 performance. In Model 1, baseline WM performance was the independent variable. In Model 2, we added HFnu during SWS. Model 1 was significant (F_(1,27)_= 70.26, p<.001; adj R^2^ =. 712), suggesting that individual difference at baseline has a strong impact on session 2 performance. Model 2 also significantly predicted performance (F_(2,26)_= 42.71, p<.001; adj R^2^ =. 749) with HFnu as a significant predictor (p=. 035). Comparing Model 1 and 2 using a likelihood ratio test, we found that Model 2 was a better fit compared to Model 1 (p=.025). In summary, while baseline WM performance provides large amount of shared variance with session 2 WM performance, HFnu during SWS added significantly more explained variation on WM performance.

## 4. Discussion

Our health is maintained through variability that allows our biological system to adjust its resources to match specific situational demands. Heart rate variability (HRV) reflects an individual’s ability to adapt the autonomic nervous system to moment-to-moment changes in her environment (Thayer & Lane, 2009). HRV has been associated with both cognitive and health outcomes, as well as linked to age-related decreases in physiological functioning. Although sleep has been shown to modulate HRV (Baharav et al., 1995; Cellini et al., 2016; Trinder et al., 2001; Whitehurst et al., 2018), the impact of this modulation has not been examined in the context of sleep related WM gains. In the current study, we investigated the functional consequence for WM of fluctuations in autonomic activity during a daytime nap in healthy young adults. We replicated the previously reported increase in parasympathetic activity in NREM sleep during daytime naps (Cellini et al., 2016). Additionally, participants showed a sleep-dependent boost in WM after sleep, and HRV during sleep was associated with WM improvement. In summary, our results provide evidence of an important role of parasympathetic activity during sleep in WM improvement in healthy young adults.

### 4.1. Nap HRV

Similar to previous nap (Cellini et al., 2016) and nighttime sleep studies (Whitehurst et al., 2018), we found vagally-mediated parasympathetic activity increased from waking to NREM sleep. These changes suggest a shift of the ANS from sympathetic to parasympathetic regulation in the transition from wakefulness to sleep. Given the parasympathetic dominance during sleep compared with wake, nighttime sleep has been described as a “cardiovascular holiday” (Trinder, Waloszek, Woods, & Jordan, 2012). Overall autonomic balance between parasympathetic and sympathetic branches is beneficial for health and cognition, whereas autonomic imbalance, indexed by low HRV and elevated sympathetic activity is associated with increased morbidity and various pathological conditions, such as cardiovascular disease, diabetes, and Alzheimer’s disease. Moreover, increased parasympathetic activity has been shown to reduce proinflammatory cytokines, and sympathetic hyperactivity is associated with increased proinflammatory cytokine production (Jarczok et al., 2015). With older adults, parasympathetic activity is less modulated during sleep compared with young adults, and no significant sleep-stage dependent variations are reported (Brandenberger et al., 2003). This profile reveals a tendency for increased sympathetic arousal and a predominant loss of parasympathetic activity in aging, which may be related to the increased number of awakenings during sleep and lower duration of SWS in this age group. The current findings in young adults of parasympathetic enhancement during sleep, and its role in WM improvement have implications for potential translational treatment strategies that target parasympathetic activity during sleep, and also suggest future studies examining the impact of age-related changes in ANS profiles on cognition.

### 4.2. Nap and Working Memory: The Functional Roles of Cardiac Activities

While a large body of studies has demonstrated the negative impact of sleep loss preceding WM performance (Choo, Lee, Venkatraman, Sheu, & Chee, 2005; Goel et al., 2009; Pasula et al., 2018), studies into the effect of post-training sleep on WM improvement are few, and none have examined this question in the context of ANS activity. We show that, compared with wake, WM improves after a daytime nap, similar to prior studies using a nap (Lau et al., 2015) and nocturnal sleep (Kuriyama, Mishima, Suzuki, Aritake, & Uchiyama, 2008b; Zinke et al., 2018), suggesting that sleep might provide an optimal brain state that facilitate WM training. We did not, however, find a significant correlation between waking HRV (as assessed with RMSSD, HF HRV, and HF_nu_) and WM in our sample. Although prior reports of HRV and WM have reported that people with higher vagally-mediated HRV perform better on WM tasks, these results were based upon median splits of the data on HRV or WM performance (Giuliano, Gatzke-Kopp, Roos, & Skowron, 2017; Hansen, Johnsen, & Thayer, 2003; Laborde, Furley, & Schempp, 2015; Spangler & Friedman, 2017). Direct correlations between waking HF HRV and WM yielded mixed results, with one study reporting a moderate correlation (Laborde et al., 2015), another yielding a borderline correlation (Hansen et al., 2003), and one showing no significant relation (Giuliano et al., 2017). In the current study, we did not find significant correlations between waking HRV and WM baseline performance or WM improvement, but instead showed a consistent pattern of an association between HRV during SWS and WM, where greater parasympathetic activity was associated with better WM improvement. Given these results, SWS, a period of naturally high levels of parasympathetic tone, should also be considered as a viable outcome measure of cardiac autonomic activity. Furthermore, parasympathetic activity during SWS, a potential biomarker of successful WM training, needs to be further studied to understand the neurophysiological mechanisms of sleep-related WM gains.

### 4.3. Working Memory Model: Slow Wave Sleep, Parasympathetic Activity and Prefrontal Functioning

We propose a conceptual model (Figure 5), that illustrates the interaction between sleep, autonomic activity, and prefrontal brain function that together and independently contribute to WM processing. We build this model on the following corpus of findings. First, studies in healthy young adults have established that people with higher waking HRV show better executive function (including WM), which is a set of cognitive abilities strongly supported by the prefrontal cortex (Thayer, Hansen, Saus-Rose, & Johnsen, 2009). This brain region is implicated in top-down control of the vagus nerve (Shaffer et al., 2014), and prefrontal cortical thickness is positively associated with vagally-mediated HRV during wake in both young and older adults (Yoo et al., 2018). Additionally, SWS and vagal activity are highly associated in both young and older adults (Brandenberger et al., 2003). Furthermore, the current study demonstrated the importance of sleep HRV for WM in healthy young adults. Taken together, prefrontal brain functioning, SWS, and parasympathetic activity, might together support sleep-related WM improvement. Aging is characterized by a decline in executive functions (Kirova, Bays, & Lagalwar, 2015), prefrontal brain atrophy (Mander et al., 2013; Salat et al., 2004), impaired sleep (Mander, Winer, & Walker, 2017), as well as decreased vagal tone (O’Brien, O’Hare, & Corrall, 1986). Though studies have established that aging is accompanied by a decline in vagally-mediated HRV during wake (De Meersman & Stein, 2007), little is known about age-related changes in HRV during sleep. Thus, it is unclear how the impact of aging on: 1) prefrontal function, 2) sleep, and 3) parasympathetic activity may be potential mediators of WM training. It remains to be seen whether the loss of parasympathetic activity during NREM sleep in older adults may mediate decreases in executive function, and/or recruitment of different brain areas to compensate for prefrontal loss. Future studies comparing younger and older populations with simultaneous brain imaging, EEG and ECG during WM training and sleep will be the next steps to further elucidate this complex interaction.

**Figure 5.**
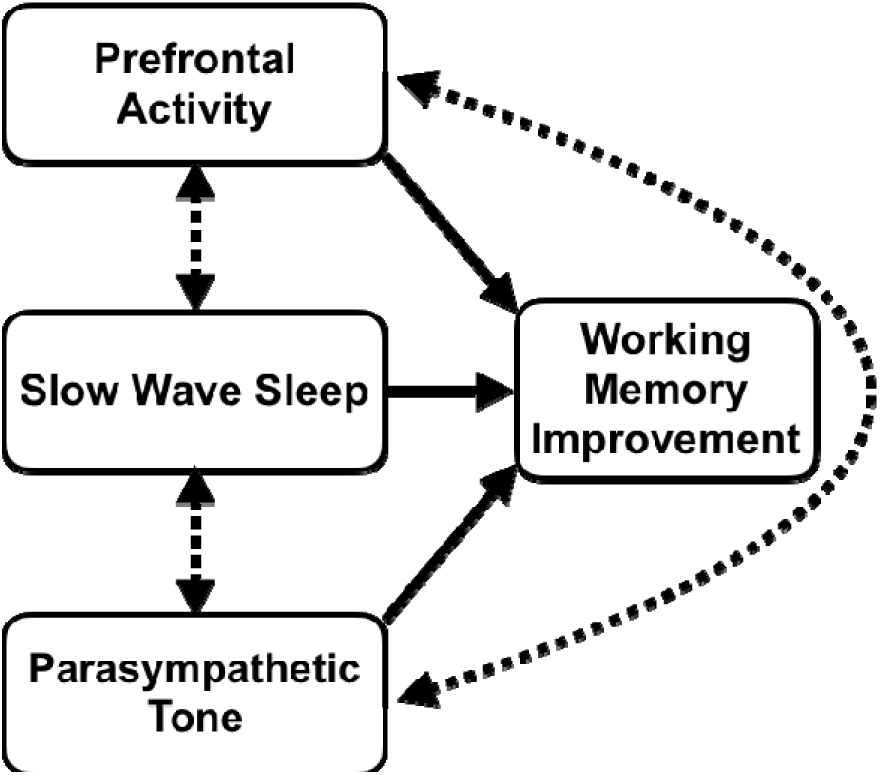
Conceptual Model: Here, we illustrate the interaction between three biological markers, Prefrontal Brain Activity, Slow Wave Sleep, and Parasympathetic Tone, which together and/or independently lead to working memory improvement in young adults.

### 4.5. Limitations

The current study has several limitations that need to be addressed. First, one limitation of this study is the reduction in subject numbers due to HRV methodological constraints. Specifically, standard practice for HRV analyses (Malik et al., 1996) requires assessment of HRV over a five-minute period of a consistent sleep stage. Due to the large amount of sleep transitions present in a daytime nap, this method decreased the number of available subjects and may have biased the sample towards less fragmented sleepers. In order to retain more statistical power, future studies should confirm these results in HRV assessments during nocturnal sleep. On the same topic, this limitation in data may underpower certain statistical comparisons, in particular for nap designs that have limited sleep compared with full night designs. For example, the current results found a significant association between performance and HFnu only, whereas the other markers of parasympathetic activity, though in a similar positive direction, did not reach statistical significance. This discrepancy may have been due to low power within certain sleep stages, and should be further investigated in a nighttime sleep study, which provides longer bouts of deep sleep.

### 4.6. Conclusion

The present study investigated the role of sleep HRV during a daytime nap in WM performance across a day. Our results confirmed that sleep benefited WM. Moreover, we showed the first evidence that the autonomic activity during sleep, but not wake, played a crucial role in WM. Thus, for heathy young adults, a daytime nap can serve as a “mini cardiovascular break” that benefits executive functions.

## Contributions

P.C. and L.N.W. contributed equally to this work; L.N.W. performed research; P.C. analyzed data; and P.C. wrote the paper.

